# Functional consequences of reductive protein evolution in a minimal eukaryotic genome

**DOI:** 10.1101/2023.12.31.573788

**Authors:** Jason Jiang, Rui Qu, Maria Grigorescu, Winnie Zhao, Aaron W. Reinke

## Abstract

Microsporidia are parasites with the smallest known eukaryotic genomes. The extent of protein loss in these organisms has been well documented, but much less is known about how compaction of microsporidia proteins affects their function. Taking a comparative genomic approach, we identified microsporidia orthologs of budding yeast proteins and show that these orthologs are enriched for essential yeast genes. We show that the median microsporidia protein is 21% shorter than its yeast counterpart and although extensive protein loss occurred after the divergence of microsporidia, reduced protein sizes were already present in microsporidian relatives. Microsporidia proteins are shorter through reduced domain lengths, diminished linker lengths, and domain loss, with 21% of microsporidia orthologs having lost domains present in yeast. On average, 34% of microsporidia orthologs have lost C-terminal residues essential for function in yeast, including 13 essential domains lost per genome. We also found that microsporidia display distinct phylogenetic patterns of domain loss, with losses occurring in a clade-specific manner. To investigate conservation of function, we used yeast complementation assays to test orthologs from several microsporidia species and their relative *Rozella allomycis*. These experiments reveal that most microsporidia proteins cannot complement their yeast orthologs, the ability to complement is about three-fold less than observed for *R. allomycis* orthologs, and proteins that do not complement are more reduced in length than their yeast orthologs. Altogether, our results demonstrate the drastic reduction of microsporidia proteins and show that these reductions have resulted in functional divergence from their fungal ancestors.

## Introduction

The change from a free-living to a host-associated lifestyle is linked with genomic reduction and loss of protein-coding genes in bacteria, archaea, protists, and animals (Moran 2002; Lind et al. 2018; Mathur et al. 2019; Slyusarev et al. 2020). Microsporidia are an extreme example of genomic minimalization. These eukaryotic intracellular parasites are related to fungi and infect a wide range of animals and protists (Keeling 2009; Murareanu et al. 2021; Bojko et al. 2022). The microsporidia *Encephalitozoon cuniculi* was the first eukaryotic parasite sequenced, revealing a 2.9 megabase genome that codes for only ∼2000 proteins, compared to ∼6000 proteins in budding yeast (Katinka et al. 2001; Giaever and Nislow 2014). Microsporidia are highly reliant on their hosts and have undergone extensive gene loss, including in many metabolic genes (Keeling and Fast 2002; Nakjang et al. 2013; Wadi and Reinke 2020; Wadi et al. 2023). In addition to protein loss, the proteins themselves are on average ∼15% shorter than their *Saccharomyces cerevisiae* orthologs (Katinka et al. 2001; Desjardins et al. 2015). Ribosomal structures from microsporidia have revealed the smallest eukaryotic ribosomes which show extreme levels of protein compaction, often with losses occurring at the N- or C-terminus (Barandun et al. 2019; Ehrenbolger et al. 2020; Nicholson et al. 2022). The proteosome has also been shown to have lost regulatory proteins and to have undergone compaction (Jespersen et al. 2022). Although the functional consequences of microsporidia protein compaction are mostly not known, it was speculated that these shorter proteins would result in reduced protein-protein interactions (Katinka et al. 2001). One example of protein reduction is the microsporidian leucine aminoacyl-tRNA synthetase which lacks a functional proofreading domain and protein synthesis at leucine residues have a high error rate (Melnikov et al. 2018).

The length and structure of proteins are constrained by evolutionary pressures. Proteins consist of independently folded domains which are the functional units of proteins, flanked by regions of amino acids known as linkers. Conservation of domain architecture has been observed across eukaryotic orthologs (Forslund et al. 2011). Proteins are shorter in prokaryotes than eukaryotes, largely explained by longer linker regions in eukaryotes as the domain sizes between the two kingdoms is similar (Brocchieri and Karlin 2005; Wang et al. 2005; Wang et al. 2011). Conserved proteins tend to be longer and longer proteins have more protein-protein interactions, suggesting a link between connectivity and protein length (Lipman et al. 2002; Warringer and Blomberg 2006). In eukaryotes, there is a slight tendency for smaller genomes to have larger average protein sizes, though microsporidia are an exception (Tiessen et al. 2012). There are fewer conserved protein domains in microsporidia than there are with other fungal species (Barrera et al. 2014). There are some examples of domain loss in microsporidia orthologs, but a comprehensive and systematic analysis of the changes that result in microsporidia protein compaction has not yet been performed (Heinz et al. 2012; Melnikov et al. 2018).

A powerful approach to explore if protein function is conserved between orthologs is to experimentally test if a diverged ortholog can replace the function of its counterpart. Studies using this approach have been facilitated by complete heterozygous knockout (hetKO) and temperature-sensitive collections in the ∼1000 essential genes in the yeast *S. cerevisiae* (Giaever et al. 2002; Li et al. 2011; Giaever and Nislow 2014; Kofoed et al. 2015). Using these collections, the ability of human genes to replace deletions in essential orthologs has been systematically measured. About half of the ∼400 human orthologs tested could rescue growth in their corresponding mutants. This result was only partly dependant on gene sequence identity and genes that function in the same process tended to display similar levels of replaceability (Kachroo et al. 2015). These types of experiments have demonstrated that conservation of protein function often extends across kingdoms, with 61% of *Escherichia coli* orthologs being able to complement yeast mutants (Kachroo et al. 2017). The ability of orthologs from other species of yeast to complement *S. cerevisiae* deletions ranges from ∼60-90% (Lai et al. 2023).

Yeast complementation has also been used to test the ability of microsporidia proteins to complement their yeast orthologs. Proteins from several species of microsporidia, including *E. cuniculi*, *Antonospora locustae*, and *Trachipleistophora hominis*, have been shown to rescue mutants in transcription, translation, the secretory pathway, and iron-sulphur cluster assembly (Hausmann et al. 2002; Hausmann et al. 2004; Slamovits et al. 2006; Upadhya et al. 2006; Goldberg et al. 2008; Freibert et al. 2017). There are also several examples of microsporidia proteins being unable to complement processes such as iron-sulphur cluster assembly or mitochondrial import (Goldberg et al. 2008; Waller et al. 2009; Freibert et al. 2017). As microsporidia genomes contain some of the smallest eukaryotic proteins, there has been interest in using complementation to understand how these minimal eukaryotic proteins function, but this approach has so far not been carried out systematically (Texier et al. 2013).

Recent ecological and sequencing efforts have identified relatives of the microsporidia (Wadi and Reinke 2020; Corsaro 2022). The closest known sister species to microsporidia is *Rozella allomycis,* which is a member of the Cryptomycota (James et al. 2013). The next closely related group of species are the early diverging microsporidia (also known as short-branched microsporidia) which are evolutionarily between *R. allomycis* and the canonical microsporidia (Bass et al. 2018). There are also the Metchnikovellids which are hyperparasites of gregarines and are basal to the canonical microsporidia (Galindo et al. 2018). Together these different microsporidian relatives provide an opportunity to understand the evolution of protein compaction in these highly reduced parasites.

A conserved microsporidia protein may be under selection to maintain the same functional ability as its budding yeast counterpart. Alternatively, the conserved microsporidia protein may be performing a different role or the context that the protein performs in has changed and thus the functional constraints the protein evolves under are different than in yeast. To distinguish between these possibilities, we used both bioinformatic and experimental approaches to study microsporidia protein evolution. We identified single-copy orthologs between microsporidia species and yeast. This analysis revealed a set of mostly essential orthologs with diminished proteins sizes and many of the lost residues are functional in their yeast orthologs. Moreover, domain loss was highly prevalent, occurring in 21% of microsporidia orthologs, with many of these lost domains essential in yeast. Additionally, we observed that domain losses occurred in a clade-specific manners in microsporidia. Using yeast complementation assays, we directly tested the function of microsporidia orthologs, observing that microsporidia proteins are significantly less likely to complement than their sister group *R. allomycis*. Our study suggests that microsporidia proteins have undergone extensive loss of function compared to their yeast orthologs and provides a system for further understanding protein evolution in a minimal eukaryotic genome.

## Results

### Microsporidia share a predominantly essential and reduced in length set of single-copy orthologs with S. cerevisiae

To compare the evolutionary properties of microsporidia proteins, we first identified orthologs shared between microsporidia species and the yeast *S. cerevisiae* using OrthoFinder (Emms and Kelly 2019). We analyzed 41 microsporidia species which we refer to as the canonical microsporidia, two species of the Metchnikovellids (Mikhailov et al. 2017; Bass et al. 2018; Galindo et al. 2018; Wadi and Reinke 2020), the early diverging microsporidia species *Paramicrosporidium saccamoebae* and *Mitosporidium daphniae* (James et al. 2013; Haag et al. 2014; Quandt et al. 2017), and the Cryptomycota species *R. allomycis*. For comparison, we also included six eukaryotic outgroup species (Supplementary table S1). We only considered single-copy orthologs as paralogous gene relationships display divergent function (Forslund et al. 2011; Soria et al. 2014) (fig. 1A and methods). On average, microsporidia species shared 438 single-copy orthologs with yeast (Supplementary fig. S1 and Supplementary table S2). The majority (61%) of these conserved microsporidia proteins are orthologous to essential yeast genes. In comparison, *R. allomycis* contained over 1000 orthologous genes with ∼40% of them being essential in yeast (Supplementary fig. S1 and Supplementary table S2).

**FIG. 1.**
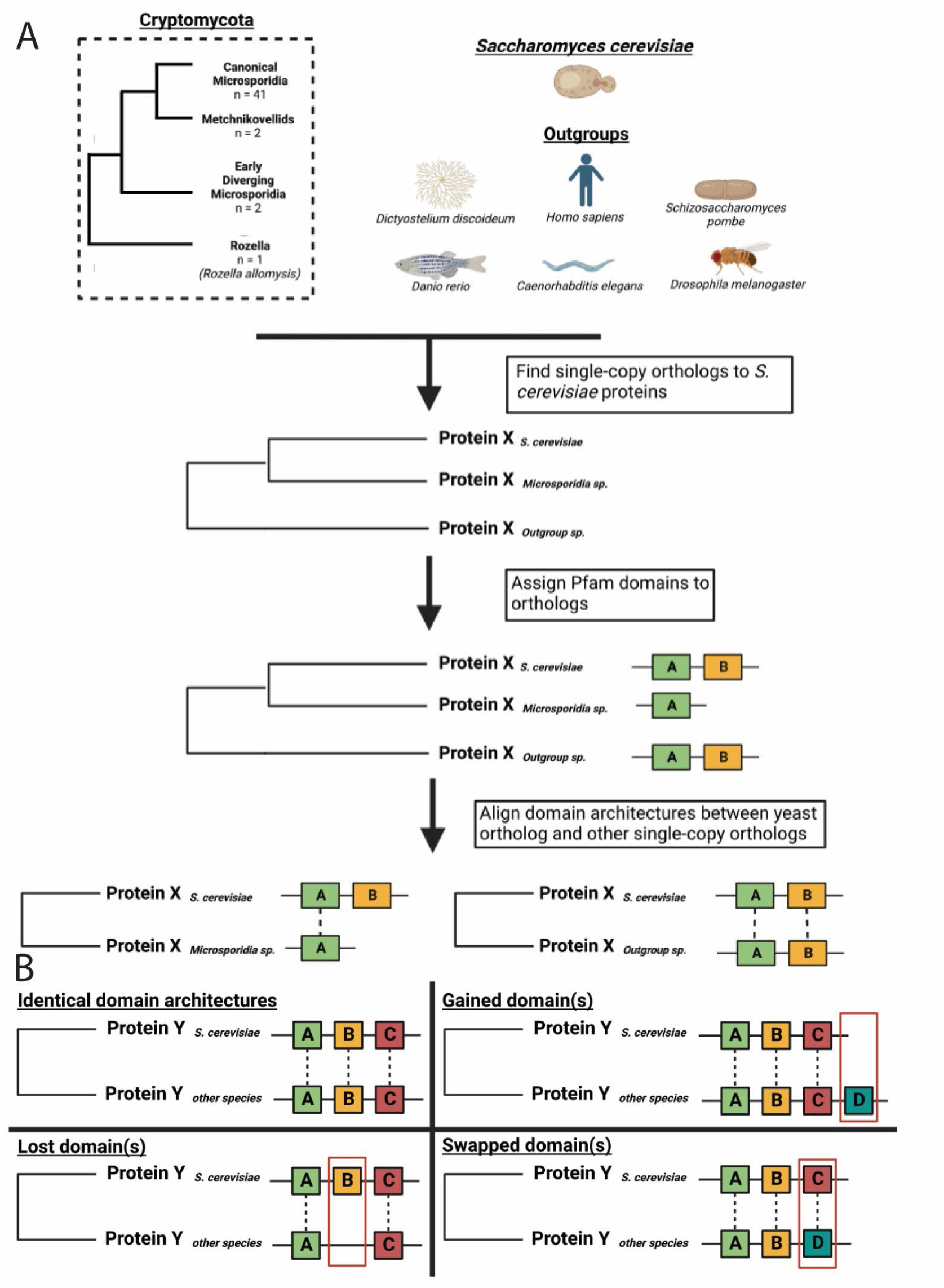
Schematic overview of identifying single-copy orthologs and aligning orthologous domain architectures. (A) Computational workflow for identification of single-copy orthologs shared between *S. cerevisiae* and microsporidian or outgroup species. Shared single-copy orthologs are identified using Orthofinder and then Pfam domains are assigned. These domain architectures are then aligned. (B) Once domains are aligned, they are then compared to identify cases where domains are either conserved, gained in the other species compared to yeast, lost in the other species compared to yeast, or swapped, with a different domain in the other species compared to yeast.

Previous reports on microsporidia genomes have reported that microsporidia proteins are shorter than their yeast orthologs (Katinka et al. 2001; Desjardins et al. 2015). To see how widely conserved this phenomenon is, we compared the length of each microsporidian ortholog to its yeast counterpart. We find that the median canonical microsporidia ortholog is 21% shorter than its yeast counterpart, compared to the median eukaryotic outgroup protein being less than 2% shorter (fig. 2A and Supplementary table S2). In both microsporidia and the outgroup, orthologs of non-essential proteins are slightly shorter than those of essential proteins (fig. 2C). Most microsporidia proteins (73%) are less than 90% the length of their yeast counterparts. In contrast, only a quarter of the outgroup orthologs are this reduced (Supplementary fig. 2). *R. allomycis* and the early diverging microsporidia represent an intermediate, as slightly over half their orthologs are less than 90% the length of their yeast counterparts (fig. 2A and Supplementary fig. 2). Microsporidia orthologs are also diverged in sequence content from their yeast orthologs, sharing only 36% sequence identity on average. The other groups are also diverged, between 38-40% sequence identity. For all groups examined, essential orthologs have a slightly higher average sequence identity than non-essential orthologs (Supplementary fig. S3A and table S2). Although microsporidia orthologs are both reduced in length and diverged in sequence identity, there is poor correlation between these two attributes (Supplementary fig. S3B).

**Fig. 2.**
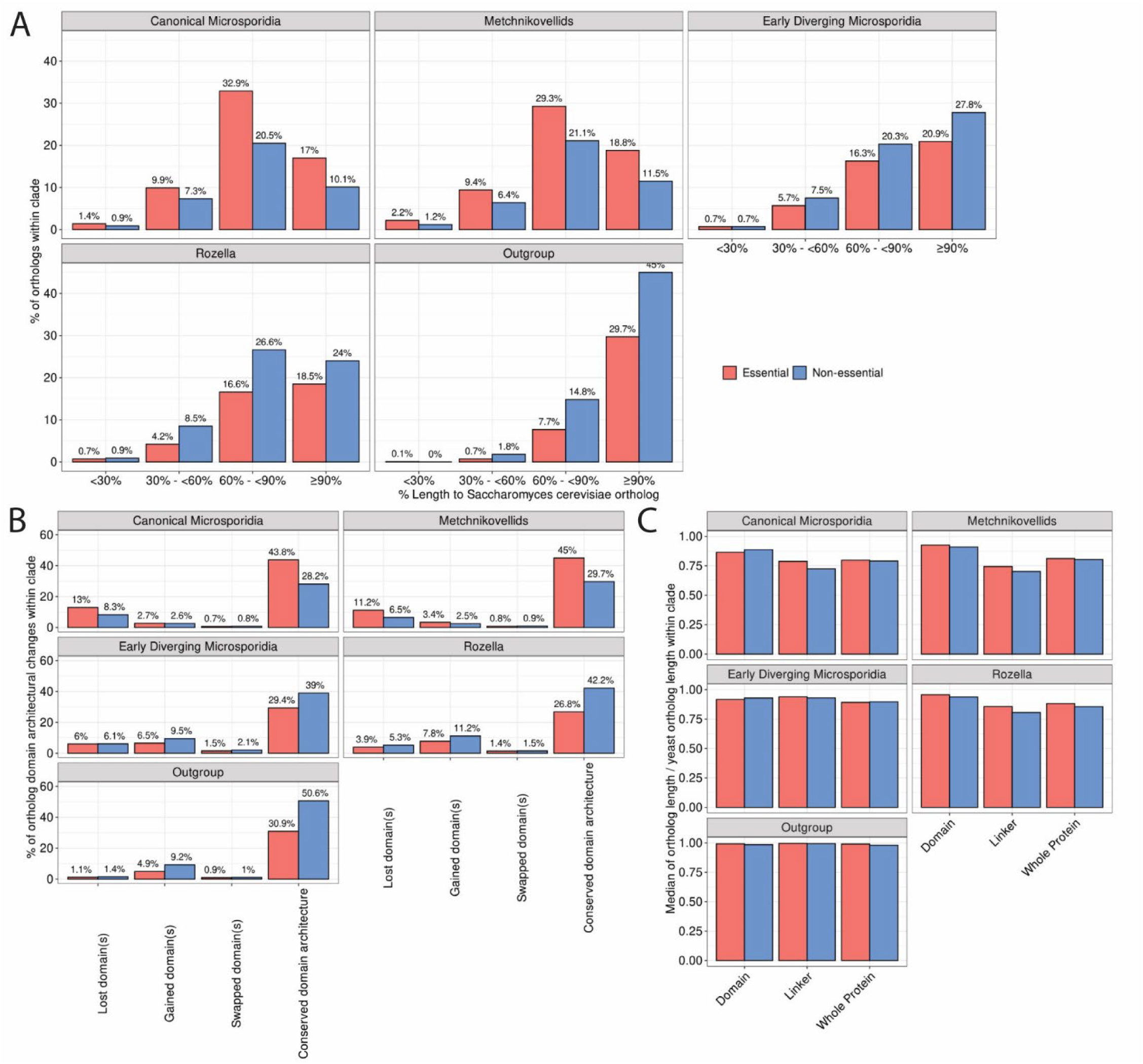
Microsporidia proteins are shorter and have lost domains compared to their yeast orthologs. (A-C) Comparison of ortholog properties of the indicated groups of species stratified by yeast-essential (red) and non-essential (blue) orthologs. (A) Distribution of protein length relative to yeast ortholog. (B) Number of indicated domain architectural change events. Orthologs with multiple domain architecture change events (ex: domain loss + domain swap) are counted toward all categories. (C) Relative length of full-length protein, domain, and linker, relative to yeast ortholog.

### Conserved microsporidia proteins have undergone widespread domain loss

To better understand how protein reduction affected the structure of microsporidia proteins, we compared domain architectures between microsporidia and yeast single-copy orthologs. Three major domain architectural changes were identified: domain loss, domain gain, and domain swap (fig. 1B and supplementary table S3). Domain architecture between most ortholog pairs was conserved, with 72% of microsporidia orthologs having identical domain architectures to their yeast orthologs (fig. 2B). *R. allomycis* and early-diverging microsporidia orthologs were conserved to a similar extent (69%) and the eukaryotic outgroup orthologs were the most conserved (82%). In canonical microsporidia, domain loss was the most prevalent architectural change, occurring in 21% of orthologs, which is about twice as much in *R. allomycis* and the early diverging species. Outgroup orthologs showed a domain loss rate of only 2.5% (supplementary table S3). We also observed higher rates of gained domains in *R. allomycis*, early diverging microsporidians, and the outgroup compared to either the canonical microsporidia or the metchnikovellids. Domain architecture changes occurred to similar extents in both essential and non-essential yeast orthologs (fig. 2B).

In addition to domain loss, microsporidia orthologs could become shorter through length reduction of their domains, linker sequences, or both. To ascertain which of these possibilities has occurred, we examined microsporidia orthologs with a conserved domain structure. Microsporidia domains have a median of 89% of their corresponding yeast ortholog domain lengths. An even greater decrease was seen in linker lengths with the median microsporidian linkers being 70% of their corresponding yeast ortholog linker lengths. In both cases, only small differences were observed between essential and non-essential orthologs. Outgroup ortholog domain and linker lengths were less reduced than in microsporidia, *R. allomycis,* and early-diverging microsporidia. Diverging species were intermediary (fig. 2C and Supplementary table S4).

### Shortened microsporidia orthologs have lost residues essential for function in yeast

Although microsporidia proteins have been observed to be shorter than their yeast orthologs, the functional consequences of these losses are not known. One possibility is that microsporidia proteins are just losing residues that are non-essential in yeast. To determine if microsporidia orthologs have lost residues that are essential in their yeast counterparts, we analyzed data from a previous study that used CRISPR-Cas9 to insert premature stop codons (PTCs) into essential yeast genes (Sadhu et al., 2018). We calculated whether the residues that were lost at the C-terminus in microsporidia orthologs were either essential, non-essential, or ambiguous in yeast (fig. 3A and Supplementary table S5). We observed that the number of essential residues lost in microsporidia orthologs was higher than in the outgroup species (fig. 3B). Of the proteins that are shorter than their yeast orthologs, 37% have lost essential residues compared to 22% of outgroup proteins (fig. 3C). An even more striking difference is observed if all proteins are considered; 34% of all microsporidia have lost essential residues compared to 12% of outgroup proteins. In addition to losing essential residues, the median microsporidia ortholog has lost all dispensable residues, with only ∼70% of outgroup orthologs having done so (fig. 3D). Together these results suggest that residues necessary for function in yeast have been lost from the C-terminus of microsporidia orthologs.

**FIG. 3.**
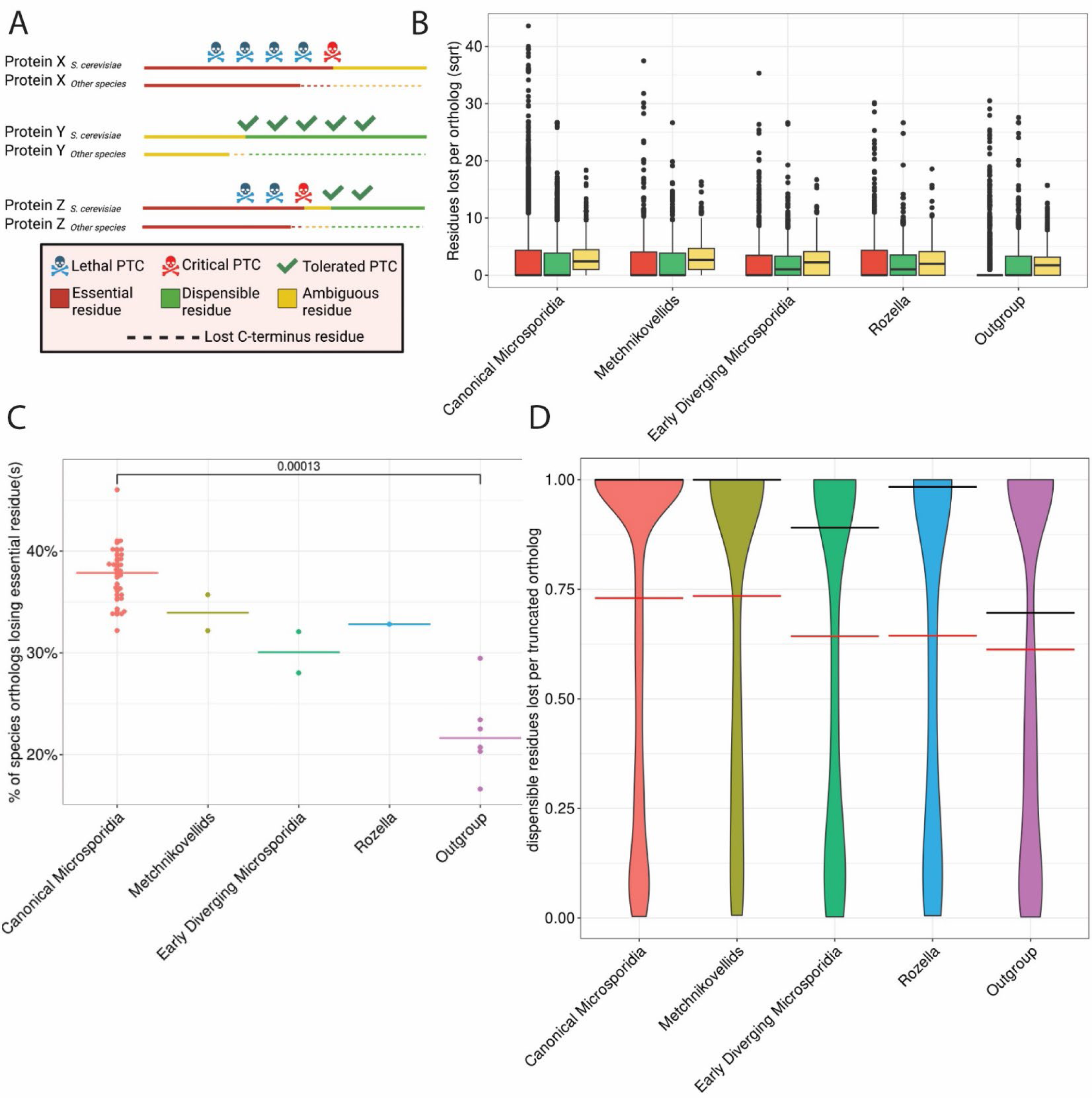
Microsporidia orthologs have lost amino acids that are essential in yeast. (A) Cartoon schematic for amino acid essentiality classifications in essential yeast genes, using premature stop codon (PTC) tolerance data (Sadhu et al. 2018). Based on a PTC being either lethal or tolerated, residues were classified as dispensable, essential, or ambiguous. (B) Box-plot showing distributions of lost amino acid types in yeast-essential orthologs for each species group. (C) Percentage of proteins from each species that are shorter than their yeast ortholog that have lost any amino acids classified as essential. Each species is a data point and the average between different species groups is shown. P-value determined using Mann-Whitney test. (D) Distribution of the percentage of dispensable residues lost in each yeast ortholog. Data is presented as a violin plot and the median in each group is shown with a black line and the mean with a red line.

Protein domains in yeast that are lost in their microsporidia orthologs might be carrying out an essential function. Using the premature stop codon data, we classified each C-terminal domain as essential, dispensable, or ambiguous (fig. 4A and Supplementary table S6) (Sadhu et al. 2018). We observed that on average each species of microsporidia has lost 13 domains that are essential in yeast, compared to 1 domain per eukaryotic outgroup species (fig. 4B).

**FIG. 4.**
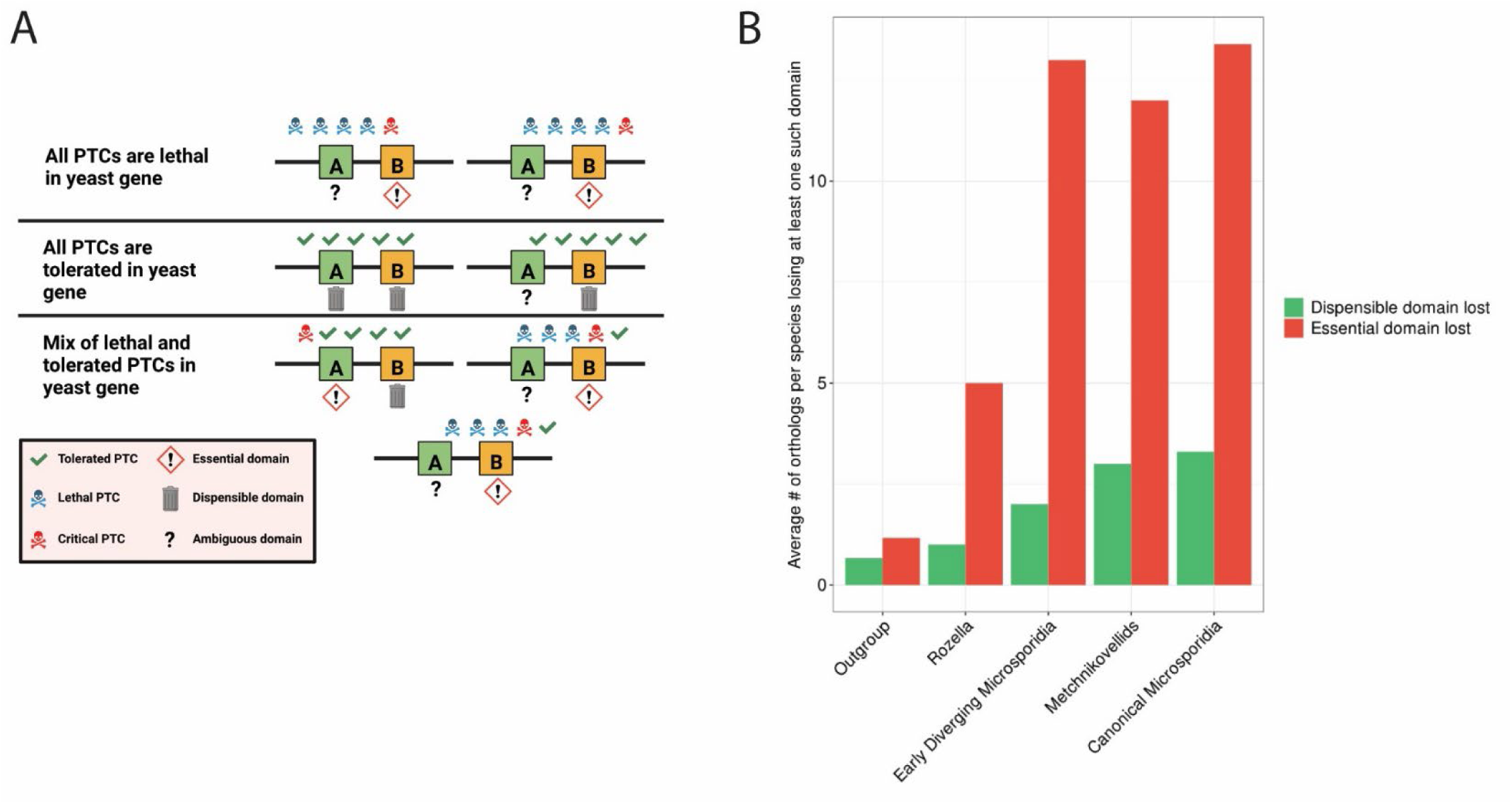
Microsporidia orthologs have lost domains that are essential in yeast. (A) Cartoon schematic of domain classification in essential yest genes with PTC tolerance (Sadhu et al. 2018). (B) Composition of lost domains in yeast-essential orthologs, displayed as different groups showing the average number of proteins that have lost domains classified as either essential or dispensable.

### Domain architectural changes in microsporidia orthologs of yeast proteins occurred early in evolution in clade-specific patterns

We then sought to determine whether domain architectural changes in microsporidia orthologs followed any phylogenetic patterns. First, we clustered orthologs by their domain architectural classifications of being conserved, lost domains, or gained/swapped domains (fig. 5A). Clustering revealed a core set of yeast genes conserved across microsporidia that have mostly conserved domain architectures. In addition, some orthologs lost in canonical microsporidia were still present in microsporidian ancestors with conserved domains. Moreover, there was a prominent cluster of genes that lost domains in most microsporidia species. However, there were also incidences of individual species and clade-restricted domain losses and gains.

**Fig. 5.**
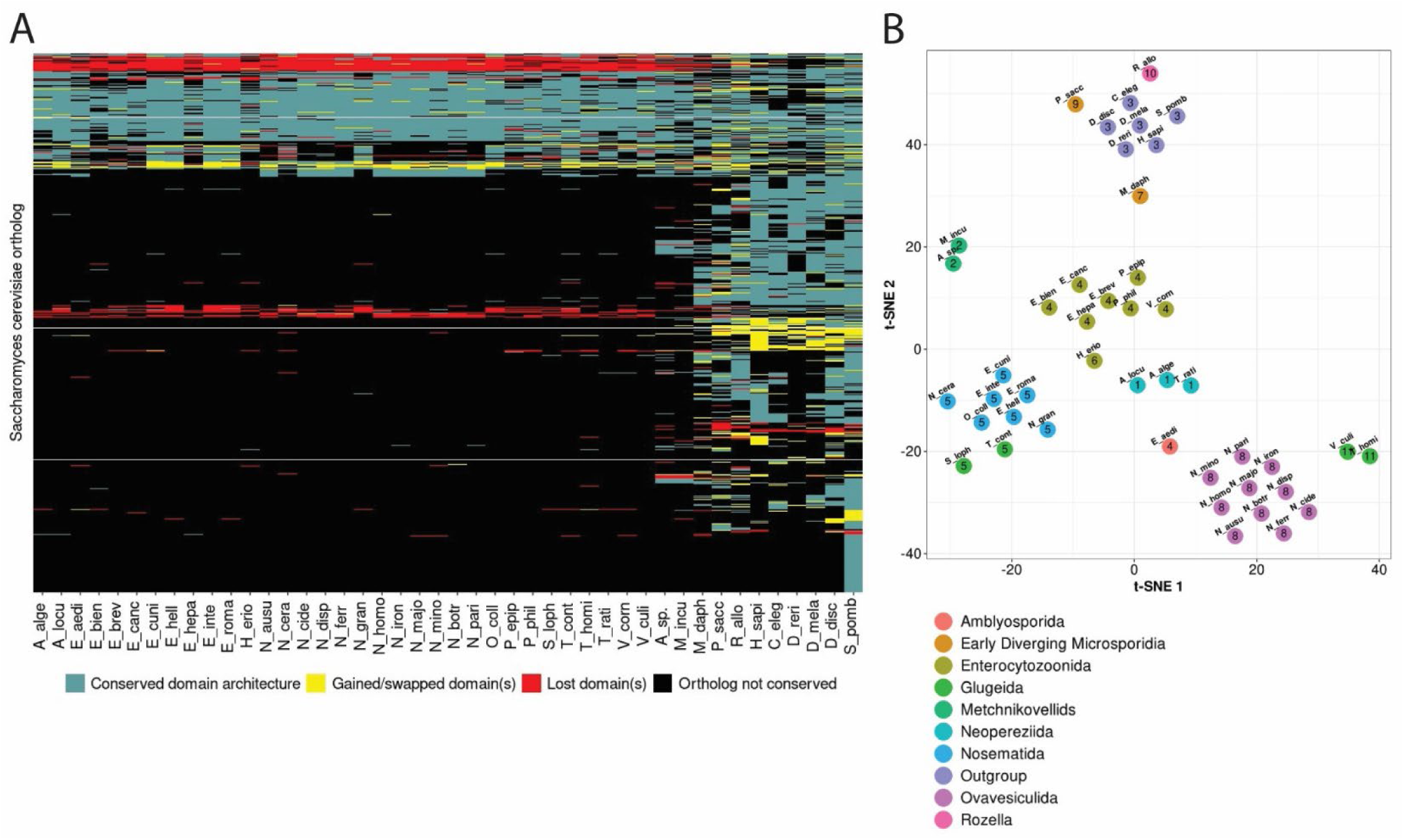
Domain architectural changes in microsporidia proteins relative to yeast occurred early in evolution, in patterns consistent with phylogeny. (A) Domain architecture conservation status of microsporidia orthologs to yeast. Rows are yeast orthologs found in at least 1 microsporidia species, and columns are orthologs from a single microsporidia species. Yeast orthologs were clustered by their similarity in domain architecture conservation patterns across microsporidia species (see methods). Species are arranged according to their broader phylogenetic clades. (B) t-SNE plot of species by composition of lost Pfam clans in orthologs. Points are colored by phylogenetic clade of species (Bojko et al. 2022) and numbered by cluster assigned through k-medoids clustering of species by lost clan content.

As groups of genes seemed to lose domains based on their phylogenetic relationships, we investigated if there were any phylogenetic patterns to the lost domains themselves. We clustered microsporidia species using lost domains in their yeast orthologs. Indeed, microsporidia have undergone patterns of domain loss consistent with their phylogenetic groupings (fig. 5B and supplementary table S7). For instance, species were separated into their broader evolutionary clades, with canonical microsporidia and microsporidia relatives grouping into distinct clusters. Moreover, we observed genera-specific patterns of domain loss, such as with *Nematocida* and *Encephalitozoon* forming their own distinct clusters.

### Most microsporidia orthologs do not complement essential yeast deletions

To experimentally determine the ability of microsporidia proteins to function in place of their *S. cerevisiae* orthologs, we used yeast complementation assays. To test if microsporidia orthologs rescue growth of essential yeast genes, we used diploid strains of yeast that contained a heterologous deletion. We transformed these strains with plasmids containing constitutive expression of each cloned ortholog that corresponded to the deletion. We then sporulated the yeast strains and performed random spore analysis by plating the resulting haploid cells on selective media to determine if the microsporidia gene could rescue the growth of the haploid deletion strain (fig. 6A). We chose representative genes and tested orthologs from three representative microsporidia species which belong to different clades (Bojko et al. 2022); *N. parisii*, *T. hominis*, and *E. cuniculi*. We also included the closest relative of microsporidia, *R. allomycis*. In total we tested 29 yeast deletions for a total of 82 tests (Supplementary table S8). We found that *R. allomycis* had the highest complementation rate, with 8/27 genes successfully complementing (30%). In contrast, complementation by microsporidia genes ranged from 8-11% and are significantly less likely to complement than *R. allomycis* (fig. 6B). Even for genes that *R. allomycis* could complement, the microsporidian complementation rate was only 33% and microsporidia complementation was only ever observed if the *R. allomycis* ortholog could also complement (Supplementary table S9).

**Fig. 6.**
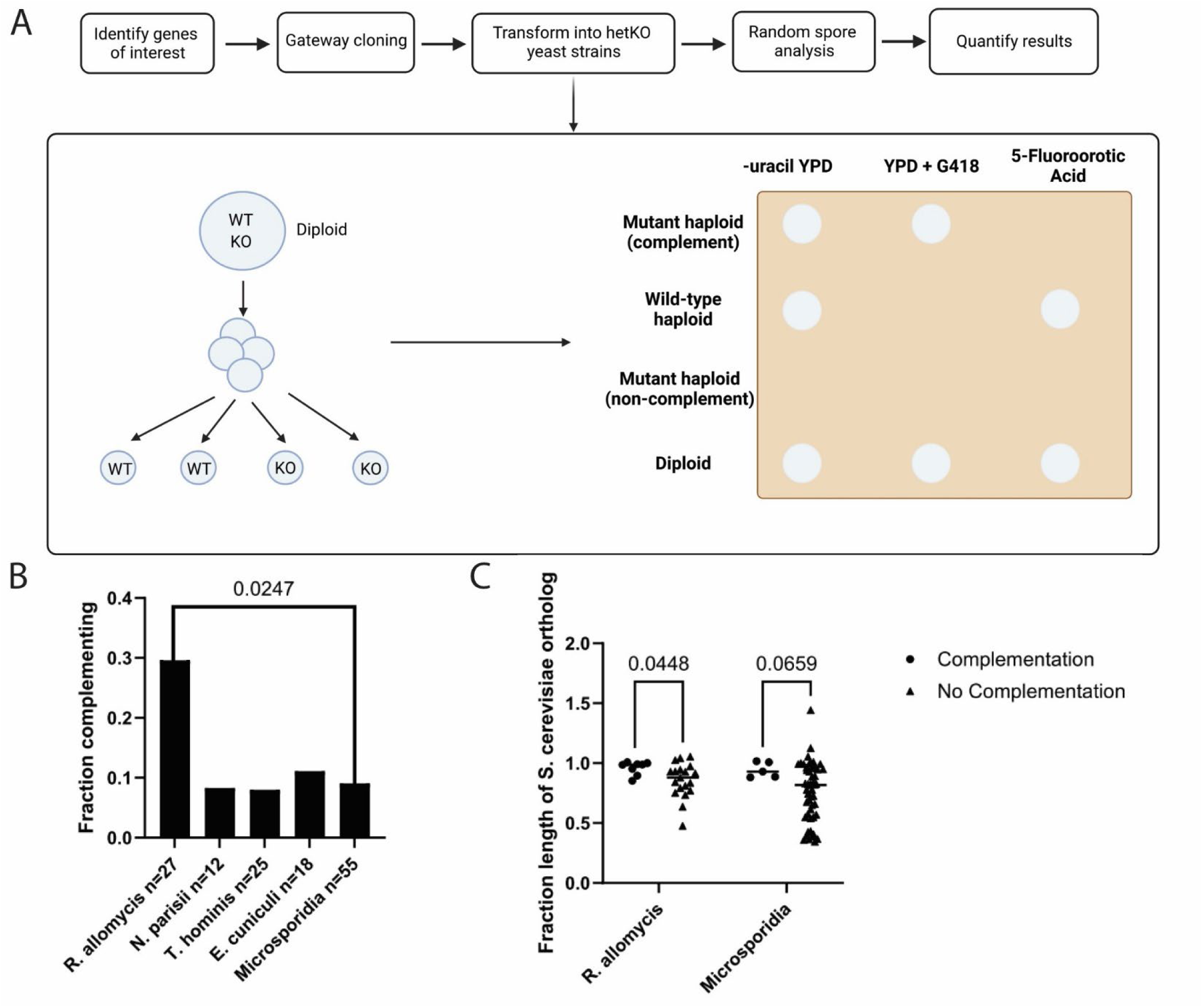
Microsporidia orthologs are less likely to complement yeast deletions than *R. allomycis* and more reduced orthologs are less likely to complement. **A.** Schematic of random spore analysis assay to determine complementation. (B) Percentage of essential yeast genes complemented by the corresponding ortholog from the indicated species. P-value was determined by two-sided Fisher’s exact test. (C) The ability of *R. allomycis* or microsporidia proteins to complement their yeast ortholog deletions compared to the length of each protein divided by length of the yeast ortholog. The p-values were determined by Mann-Whitney test.

We then compared length and domain conservation in the proteins that were functional for complementation. *R. allomycis* proteins that complemented have an average length 96% that of their yeast orthologs. In contrast, *R. allomycis* proteins that could not complement were significantly shorter with an average length 86% that of their yeast orthologs (fig. 6C). Microsporidia proteins that complemented had an average length 94% that of their yeast orthologs, while those that could not complement had an average length 76% that of their yeast orthologs (fig. 6C and Supplementary table S9). The most reduced *R. allomycis* protein that complemented was 85% the length of its yeast ortholog and the most reduced microsporidia protein that complemented was 88% the length of its yeast ortholog.

One of the reduced microsporidia proteins that complemented was the ribosomal stalk protein RPP0 for which we observed complementation for the *R. allomycis*, *E. cuniculi*, and *N. parisii* orthologs (Supplementary tables S8 and S9). The C-terminal part of the yeast RPP0 protein contains regions which are necessary for facilitating interactions with other ribosomal proteins and C-terminal truncations of RPP0 diminish yeast growth (Supplementary fig. S4A) (Santos and Ballesta 1995). These regions are missing in microsporidia orthologs. To determine if yeast RPP0 that was missing C-terminal residues could rescue to a similar extent, we generated RPP0 truncation mutants and observed that the two microsporidia, the full-length yeast, and the truncated yeast versions could all support growth by the same amount (Supplementary fig. S4B).

We then determined the reported rate of complementation of yeast deletions with microsporidia proteins by compiling a list of previously reported complementation assays (Supplementary table S10). The rate reported in published articles is 9 out of 25 (36%) proteins complementing, and in theses 1 out of 7 (14%), for an overall reported complementation rate of 31%. However, some of these proteins that complemented were cloned using exogenous mitochondrial targeting signals, and if these cases are omitted, 6/28 (21%) complement (Supplementary fig. S5A). We then compared the length of the microsporidia proteins tested to their yeast orthologs. Proteins that don’t complement are more reduced (Supplementary fig. S5B). The combined complementation data of our study and previous reported results shows that microsporidia proteins that complement are significantly less reduced (Supplementary fig. S5C). Altogether, our analysis indicates that most microsporidia proteins cannot complement deletions of their yeast orthologs and that more reduced proteins are less likely to complement.

## Discussion

Reductive evolution is a hallmark of intracellular microbes (Moran 2002). Using both computational and experimental methods we show that microsporidia proteins have undergone functional losses compared to their budding yeast counterparts. We show that microsporidia proteins are unique to other eukaryotic proteins in several ways. Microsporidia orthologs are enriched for essential genes, have undergone extreme reduction in length, and have lost residues and domains essential in yeast. The loss of functional residues and domains we observe is an underestimation, as only the impact of C-terminal residues has been determined, and it is likely that many of the residues we classify as ambiguous are necessary for function. Together our results suggest that microsporidia proteins have experienced widespread loss of the corresponding residues necessary for function of yeast proteins. One limitation of our approach is orthologs are identified based on sequence identity, which can fail to identify highly divergent microsporidia orthologs (Nakjang et al. 2013). Approaches in the future using structural based detection may be able to overcome this limitation (Mascarenhas dos Santos et al. 2022; Svedberg et al. 2023).

We observe that most microsporidia proteins cannot complement essential yeast deletions and that microsporidia proteins complement at a lower rate than has been observed for other species. Yeast complementation assays have been used on orthologs from a variety of different species with complementation rates ranging from ∼50-90% (Kachroo et al. 2015; Kachroo et al. 2017; Lai et al. 2023). Although we only tested a subset of microsporidia genes for their ability to complement yeast deletions, we provide ∼3 times more complementation tests than have been previously reported for microsporidia. The overall rate of complementation we see is somewhat lower than what has been reported in the literature, though our study includes more highly reduced proteins and is likely more representative of the ability of microsporidia proteins to rescue yeast deletions. One reason that microsporidia proteins display low complementation is not localizing correctly in yeast and adding a tag has been shown to correct localization and rescue function (Goldberg et al. 2008). Another possibility is that expression levels are too low, but many reported microsporidia complementation tests used multi-copy plasmids and strong reporters and yet didn’t observe complementation (Supplementary table S10). Another explanation for failure to complement is that an essential interaction is no longer maintained between the microsporidia ortholog and a yeast protein. It could be that microsporidia proteins have rapidly co-evolved and the microsporidia protein can’t function in the yeast context due to differences in their interacting partners. This type of evolution was recently shown for the anaphase-promoting complex, where the replacement of multiple genes from other yeast species at the same time was necessary to restore function (Lai et al. 2023). Alternatively, the microsporidia protein could have evolved to no longer carry out the essential function of its yeast ortholog. Further investigation will be necessary to understand the explanation for each orthologs’ inability to complement.

Our results suggest that microsporidia evolution has resulted in progressive reduction and loss of function compared to other eukaryotes. A loss of both conserved genes and metabolic pathways has been observed in *R. allomycis* compared to other fungi, with more losses occurring in the early diverging microsporidia species (James et al. 2013; Haag et al. 2014; Quandt et al. 2017; Wadi et al. 2023). Additional losses were reported in the metchnikovellids and canonical microsporidia, with gene loss occurring to similar extents in these groups (Mikhailov et al. 2017; Galindo et al. 2018; Wadi et al. 2023). Consistent with what has been shown for conservation of proteins, we observe that *R. allomycis* and the early diverging microsporidia are more reduced than other eukaryotes, but not to the same extent as the metchnikovellids and the canonical microsporidia. We observe similar trends for domain loss and for residues that are functional in yeast. Together, our data suggests that protein compaction and domain loss had already occurred before the divergence of the canonical microsporidia. In complementation assays, we observe that *R. allomycis* can rescue the function of deletions at a rate that is higher than microsporidia, but less than species of yeast, providing further evidence that although *R. allomycis* is a parasitic species, it represents an evolutionary intermediate between free-living fungi and microsporidia.

Although microsporidia were discovered over 160 years ago, work on defining their core functional mechanisms has largely been unexplored. Genomic approaches have been extremely useful at defining what genes are present, but there is a lack of genetic tools available to test their function (Katinka et al. 2001; Reinke and Troemel 2015; Wadi and Reinke 2020). Recent high resolution structural approaches have documented how microsporidia proteins and complexes undergo compaction (Barandun et al. 2019; Ehrenbolger et al. 2020; Jespersen et al. 2022; Nicholson et al. 2022). Biochemical and ancestral reconstruction approaches have also been extremely valuable to understanding functional evolution of proteins (Goldberg et al. 2008; Freibert et al. 2017; Dean et al. 2018). The use of yeast as a heterologous system to study these proteins provides a powerful approach to systemically determining functional conservation (Kachroo et al. 2015). Our finding that some *R. allomycis* proteins can complement yeast deletions, but not the corresponding microsporidia orthologs, provides a comparative system to further explore evolution of microsporidia protein function. Using approaches such as chimeras and determining differences such as localization and protein-protein interactions will be useful for exploring how microsporidia orthologs have functionally diverged.

## Materials and Methods

### Identification of *Saccharomyces cerevisiae* single-copy orthologs and alignment of domain architectures

Orthogroups between proteomes were identified with OrthoFinder v2.5.4 (Emms and Kelly 2019), using BLAST 2.5.0+ (Johnson et al. 2008) for the all-versus-all search step. A total of 53 proteomes were used, including microsporidia genomes, microsporidia relatives, *S. cerevisiae*, and outgroups. The proteomes used and their accession numbers are listed in Supplementary table S1. After the initial OrthoFinder-step, orthologs from 8 species were excluded for being judged to be low quality assemblies (Wadi et al. 2023). Orthogroups containing a single-copy *S. cerevisiae* ortholog and at least one single-copy ortholog from any other species were identified for downstream analyses. *S. cerevisiae* orthologs were classified as essential or non-essential using a previously published list of essential yeast genes (Kofoed et al. 2015).

Protein domains in each ortholog were assigned using hmmscan from HMMER 3.3.2 (Eddy 2011) and Pfam 35.0 (Mistry et al. 2021). Pfam family-specific bit-score cut-offs for protein domain families were used over fixed E-value thresholds, due to previously demonstrated improvement benefits in using these bit-score cutoffs over E-values (Punta et al. 2012). Overlapping domain assignments from hmmscan were resolved with cath-resolve-hits 0.16.10 (Lewis et al. 2019) to obtain the final domain architectures for each ortholog. Within each orthogroup, the *S. cerevisiae* single-copy ortholog was aligned with all other single-copy ortholog in the orthogroup. To facilitate alignment, all domains in domain architectures were first mapped back to their Pfam clans, as domain families within clans have conserved function (Forslund et al. 2011; Mistry et al. 2021). Within each domain architecture, the clans were converted to unique letter representations. Finally, the letter representations of clans within each ortholog domain architecture were aligned with the Needleman-Wunsch algorithm (Needleman and Wunsch 1970), as has been previously done for domain architectural alignments (Forslund et al. 2011), using a match score of 0, a mismatch penalty of −3, and a gap penalty of −10.

### Identification of domain architectural changes from domain architecture alignments

We defined three categories of domain architectural changes in orthologs compared to their *S. cerevisiae* orthologs: domain losses, domain gains, and domain swaps. Domain losses in species orthologs were inferred as a gap in the alignment of the species domain architecture compared to its *S. cerevisiae* ortholog. Domain gains were inferred as gaps in the aligned *S. cerevisiae* ortholog domain architecture compared to its species ortholog. Domain swaps were inferred as mismatching positions between the aligned domain architectures. For species-*S. cerevisiae* ortholog pairs where the species ortholog lacked domain assignments, we considered the species ortholog to have lost all the domains present in its *S. cerevisiae* ortholog. Conversely, if the *S. cerevisiae* ortholog lacked domain assignments, we considered the species ortholog to have gained all its domains relative to the *S. cerevisiae* ortholog. To account for erroneous missing domain classifications, we excluded *S. cerevisiae* -species ortholog pairs with domain loss where the species ortholog was not shorter than its *S. cerevisiae* ortholog by at least 85% of the length of all of its lost domains from *S. cerevisiae*. We employed this heuristic filter to exclude instances of domain loss classified by our pipeline that were due to high sequence divergence in the species ortholog sequence from *S. cerevisiae*. Such *S. cerevisiae* -species ortholog pairs were excluded from all downstream analyses.

### Comparison of domain and linker lengths between microsporidia and yeast single-copy orthologs

Total domain length for an ortholog was calculated as the summed lengths of all domains within the ortholog. For orthologs without domain assignments, the total domain length was defined as zero. We excluded ortholog pairs from orthogroups where neither the other species nor *S. cerevisiae* ortholog had domain assignments. Linker lengths were defined as the total protein length for an ortholog with total domain length subtracted. To compare total domain and linker lengths between ortholog pairs, we divided the length in the other species by the length in the *S. cerevisiae* ortholog.

### Identification and classification of lost C-terminal amino acids in single-copy yeast-essential genes

First, we classified each amino acid within all essential *S. cerevisiae* genes as either essential, dispensable or ambiguous for *S. cerevisiae* viability, using a dataset of premature stop codons (PTCs) inserted into all essential yeast genes and their effects on yeast viability (Sadhu et al. 2018). All amino acids coming at or before a yeast-lethal PTC were classified as essential. All amino acids coming at or after a tolerated PTC were classified as dispensable. However, amino acids were classified as ambiguous in the following cases: (1) amino acids coming after the C-terminus PTC in a gene and the C-terminus PTC is lethal, (2) amino acids coming before the N-terminus PTC and the N-terminus PTC is tolerated, and (3) amino acids between a lethal and tolerated PTC. Amino acids in (1) were excluded, as it is unclear whether amino acids after the C-terminus lethal PTC are essential or dispensable. Amino acids in (2) were excluded as it is unclear if essential amino acids exist upstream of the N-terminus tolerated PTC. Amino acids in (3) were excluded as it is unclear if amino acids exist upstream of the tolerated PTC, in the region between the lethal and tolerated PTC.

To identify lost C-terminal amino acids in orthologs to yeast, we first identified yeast-essential orthologs that were shorter than their yeast equivalents. Such ortholog pairs were aligned with the Smith-Waterman local alignment algorithm, using the BLOSUM45 substitution matrix for microsporidia orthologs to account for their high sequence divergence (Nakjang et al. 2013), and the BLOSUM62 matrix for all other species. Lost C-terminal residues were defined as residues in the yeast ortholog coming after the end of the local alignment and were classified as described above.

### Classification of dispensability of lost domains in single-copy orthologs to yeast

We classified domains in all essential *S. cerevisiae* genes as “essential”, “dispensable” or “ambiguous” using the PTC dataset in essential yeast genes and their effects on yeast viability (Sadhu et al. 2018). Within each essential yeast gene, we indicated the C-terminal most PTC as the “critical PTC”. Essential domains were classified as domains for which the critical PTC appears upstream of the domain end, suggesting that truncation/loss of the domain by the critical PTC led to yeast mortality. Moreover, we also classified domains with the critical PTC immediately downstream of its end as essential, to account for cases in which the real boundaries of the domain extend beyond the predicted boundaries and the domain was actually disrupted by the critical PTC. Dispensable domains were classified as domains with a dispensable PTC upstream of its start, suggesting that the domain could be completely removed without affecting yeast fitness. All other domains were classified as “ambiguous”. All lost domains in *S. cerevisiae* -essential orthologs were classified, per the classification of the domain in their corresponding *S. cerevisiae* ortholog.

### Hierarchical clustering of single-copy orthologs to yeast by domain architecture

First, a table was constructed of all single-copy yeast orthologs with at least one single-copy ortholog from another species. Columns of the table were yeast orthologs, rows were the non-yeast species (microsporidia and outgroups) and cells held information on the domain architecture conservation between the yeast ortholog and the single-copy ortholog from the species in that row. Domain architecture conservation between yeast-species ortholog pairs were classified as follows. Ortholog pairs with identical domain architectures were classified as “Conserved domain architecture”. Ortholog pairs where the species ortholog had lost one or more domain were classified as “Lost domain(s)”. Ortholog pairs having gained or swapped one or more domains were classified as “Gained/swapped domain(s)”. If a species ortholog had both lost and swapped domains, we classified it as “Lost domain(s)”. In cases where a species did not have a single-copy ortholog to a particular yeast ortholog, we classified that cell as “Ortholog not conserved”. With this table, a distance matrix was calculated for the *S. cerevisiae* orthologs, based on their dissimilarity in domain architecture conservation across species. Gower’s distance was used to calculate distances for these categorical variables, using the daisy function from the cluster v2.1.2 R package (Maechler et al. 2022). Yeast orthologs were then clustered by similarities in domain architecture conservation patterns across species, using the agnes function from the cluster R package for agglomerative hierarchical clustering with the distance matrix.

### Generation of t-SNE plot of ortholog lost domains

First, a list of all unique lost Pfam clans was collected for each species, using the aligned ortholog domain architectures for each species and their yeast orthologs. A dissimilarity matrix was constructed between each species and their set of unique clan losses using Gower’s distance, with the daisy function from R package cluster v2.1.2 (Maechler et al. 2022). Using this matrix, species were then clustered by their similarity in lost clans using k-medoids clustering, using the pam function from cluster. The optimal number of clusters (k) for k-medoids clustering was selected by clustering with k = 2 to k = number of species - 1, and selecting the k-value that maximizes silhouette width of the clustering (Rousseeuw 1987), which is a measure of how well items fit in their clusters. The clustered species were visualized using t-SNE, passing the dissimilarity matrix into the Rtsne function from the Rtsne v0.15 R package (Krijthe 2015). Maximum iterations for t-SNE was set to a high value of 5000, to ensure convergence to stable t-SNE plots. Ideal perplexity was chosen by iterating from 5 to (number of species – 1 / 3), and manually inspecting the resulting t-SNE graphs. This range of perplexities was chosen based on the recommended range of perplexities of 5 – 50 (Maaten and Hinton 2008), and from the documentation of Rtsne stating a maximum perplexity of (number of rows in data – 1 / 3).

### Cloning of microsporidia genes

Genes to be cloned were chosen using two strategies. For the first strategy we chose 14 genes where there were *R. allomycis*, *T. hominis,* and *N. parisii* orthologs. For 7 of these genes the length ratio of the *N. parisii* protein compared to its yeast ortholog had a ratio greater than 0.99. For the other 7 genes this ratio was less than 0.9. For the second strategy, 15 genes chosen at random where there were *R. allomycis, T. hominis,* and *E. cuniculi* orthologs. All genes chosen were required to encode for *R. allomycis* proteins that are less than 479 amino acids, to facilitate cloning using gene synthesis. Genes from *Nematocida parisii*, *Encephalitozoon cuniculi*, and *Saccharomyces cerevisiae* were PCR amplified from *N. parisii* ERTm1*, E. cuniculi* E3, and S. cerevisiae BY4741 genomic DNA, respectively. Using ApE (Davis and Jorgensen 2022), primers for PCR were designed for each gene of interest, with a target melting temperature of ∼55 °C. To facilitate Gateway Cloning, partial attB sequences were appended to the ends of the primers. The 5’ end of the forward primer contained the sequence ACAAAAAAGCAGGCTCA, whereas the 3’ end of the reverse primer contained the sequence GGGGACCACTTTGTACAAGAAAGCTGGGTT. Primers were designed from the start codon of every gene until the last codon before the stop. Next, a two-step PCR was conducted. In the first step, the gene of interest was amplified from genomic DNA using the gene-specific primers and Phusion polymerase. The second step amplified the product of the first step with attB primers, which appended the complete attB1 sequence to the 5’ end of the gene and attB2 sequence to the 3’ end. Amplicons from the two-step PCR were PEG-purified and successful amplification of genes was verified by agarose gel electrophoresis. Genes from *R. allomycis, T. hominis,* and some *E. cuniculi* genes were ordered as codon optimized synthesized genes from IDT. These genes were ordered with attB1 and attB2 flanking sequences.

The PCR products were inserted into the pDONR221 entry vector in the BP reaction of the Gateway cloning system and transformed into competent DH5-α *E. coli* cells to obtain entry clones. The extracted entry clones were then inserted into the pAG416GPD-+6Stop (Laurent et al. 2020) destination vector by Gateway LR reaction and transformed into competent E. coli to obtain expression clones. All entry and expression clones were verified by restriction digests and Sanger sequencing. All oligos and gene sequences are shown in Supplementary table S11.

### Transformation of expression clones into mutant yeast strains

Expression clones were transformed into yeast strains as described previously (Reece-Hoyes and Walhout 2018). The empty destination vector, pAG416-GPD-ccdB+6Stop, was also transformed into cells as a negative control. Yeast cells from transformed hetKO cells were plated on -uracil dextrose + G418 plates at 30°C for 2 - 3 days, to select for both successful transformation and presence of the G418-resistance knockout cassette. Yeast cells from transformed temperature-sensitive strains were plated onto -uracil dextrose plates and grown at 25 °C for 5 days.

### Heterozygous knockout strain complementation testing

The ability of yeast and microsporidian orthologs to complement their corresponding heterozygous-knockout yeast strains was tested with random spore analysis (protocol adapted from: https://openwetware.org/wiki/Springer_Lab:_Random_Spore_Analysis). Single colonies of transformed yeast cells were suspended in 2 mL of 1% potassium acetate, and sporulated for 5 days in a rotary shaker at 30°C. Successful sporulation was confirmed with the observation of tetrads using microscopy. Cells were washed and resuspended in 500µL milli-Q water, 25µL 5mg/mL 20T Zymolyase, and 5µL β-mercaptoethanol. The spore mixture was placed on a nutating mixer overnight at 30°C. Tetrads were disrupted the following day by the addition of Zirconium beads and shaking on a Disruptor Genie at 3000 RPM for 10 minutes. Cells were then isolated with centrifugation for 10 minutes at 10,000 RCF. To further remove any diploids, the Zymolyase treatment was repeated with 20mg/mL 20T Zymolyase, followed by Zirconium bead disruption the next day. The cells were then plated onto -uracil dextrose plates and incubated at 25°C for 5 days to select for colonies containing the plasmid of interest. Colonies were replica plated onto YPD+G418 and 5-FOA plates to test for complementation. Haploid yeast cells that complement grow on YPD+G418 but not 5-FOA, and are expected to grow in a ∼1:1 ratio with haploid yeast cells with the wild-type gene copy, which grow on 5-FOA but not YPD+G418. Cells that could grow on both plates were considered diploids. The corresponding empty vector negative control was also tested for any genes that showed complementation across all strains. Random spore analysis was repeated once with each strain, to generate at least two biological replicates.

### Testing RPP0 protein functionality via dilution assays

RPP0 truncation mutants were cloned as described above. The yeast truncation mutants, and the *N. parisii* and *E. cuniculi* orthologs were transformed into the RPP0 hetKO strain and sporulated as described above. Dilution assays were performed as described previously (Petropavlovskiy et al. 2020). For each plasmid, three complementing haploid mutant colonies were picked and grown separately overnight in -uracil dextrose medium at 25 °C, generating three different biological replicates. An empty vector negative control was picked from wild-type haploid colonies. Cell concentrations of each culture were equalized to an OD600 of 1.0 and added to the first row of a 96-well plate. Five 5X dilutions were then performed down the rows of the plate. Next, using a spotting tool, equal amounts of each sample were spotted onto a -uracil dextrose plate. Plates were then incubated for 2 days at 25 °C, 35 °C, or 37 °C. Images were taken of each plate after the 2-day incubation. Then, using ImageJ (Schneider et al. 2012) the images were adjusted so that the white colonies showed sharp contrast against the dark background. Then, the amount of growth of each sample and dilution was determined by measuring the grey value of each. The grey value was also measured for five points amongst the background, which was then averaged and subtracted from the grey value measurements for each of the samples. A mean value of each condition was then generated by averaging all the grey value measurements of the condition from each biological and technical replicate. Across each temperature, the mean values for all strains were then normalized against the value of the full-length yeast gene.

### Data and code availability

Code for analyses was written in R (R Development Core Team 2019) and Bash. Scripts were organized into a pipeline using Snakemake (Köster and Rahmann 2012). All software needed to run the pipeline were deployed with a Singularity X.Y.Z container. Code for this study was deposited to https://github.com/Jason-B-Jiang/snakemake-domain-architectures.

## Acknowledgements

We thank Qingyuan Huang, Edward James, Yin Chen Wan, and Jonathan Tersigni for providing helpful comments on the manuscript. We thank Brenda Andrews for the kind gift of yeast mutant strains. We thank Grant Brown for the kind gift of the pAG416-GPD-ccdB+6Stop vector. We thank Louis Weiss for providing *E. cuniculi* DNA. We thank Aashiq Kachroo for providing protocols and advice. We thank Alexander Ensminger, Marc Meneghini, and their labs for advice on working with budding yeast. This work was supported by a Canadian Institutes of Health Research grant no. 400784 (to A.R.), an Alfred P. Sloan Research Fellowship FG2019-12040 (to A.R.). and the Natural Sciences and Engineering Research Council of Canada Undergraduate Student Research Awards to J.J., and Undergraduate Summer Research Program awards to R.Q. and M.G.

## Supplementary figures and tables

**Supplementary table S1. List of genome assemblies used in this study.**

**Supplementary table S2. Length, essentiality, and identity of microsporidia orthologs of yeast proteins.**

**Supplementary table S3. Conservation of domain architecture.**

**Supplementary table S4. Domains and linker lengths.**

**Supplementary table S5. Comparison of PTC data of microsporidia orthologs.**

**Supplementary table S6. Classification of lost domains in yeast-essential orthologs.**

**Supplementary table S7. Identity of domains conserved or lost in yeast orthologs.**

**Supplementary table S8. Results of random spore analysis assays.**

**Supplementary table S9. Microsporidia ortholog complementation of essential yeast proteins.**

**Supplementary table S10. Previously reported results of complementation of yeast mutants with microsporidia genes.**

**Supplementary table S11. Oligos and sequences of genes used in this study.**

**Supplementary fig. S1.**
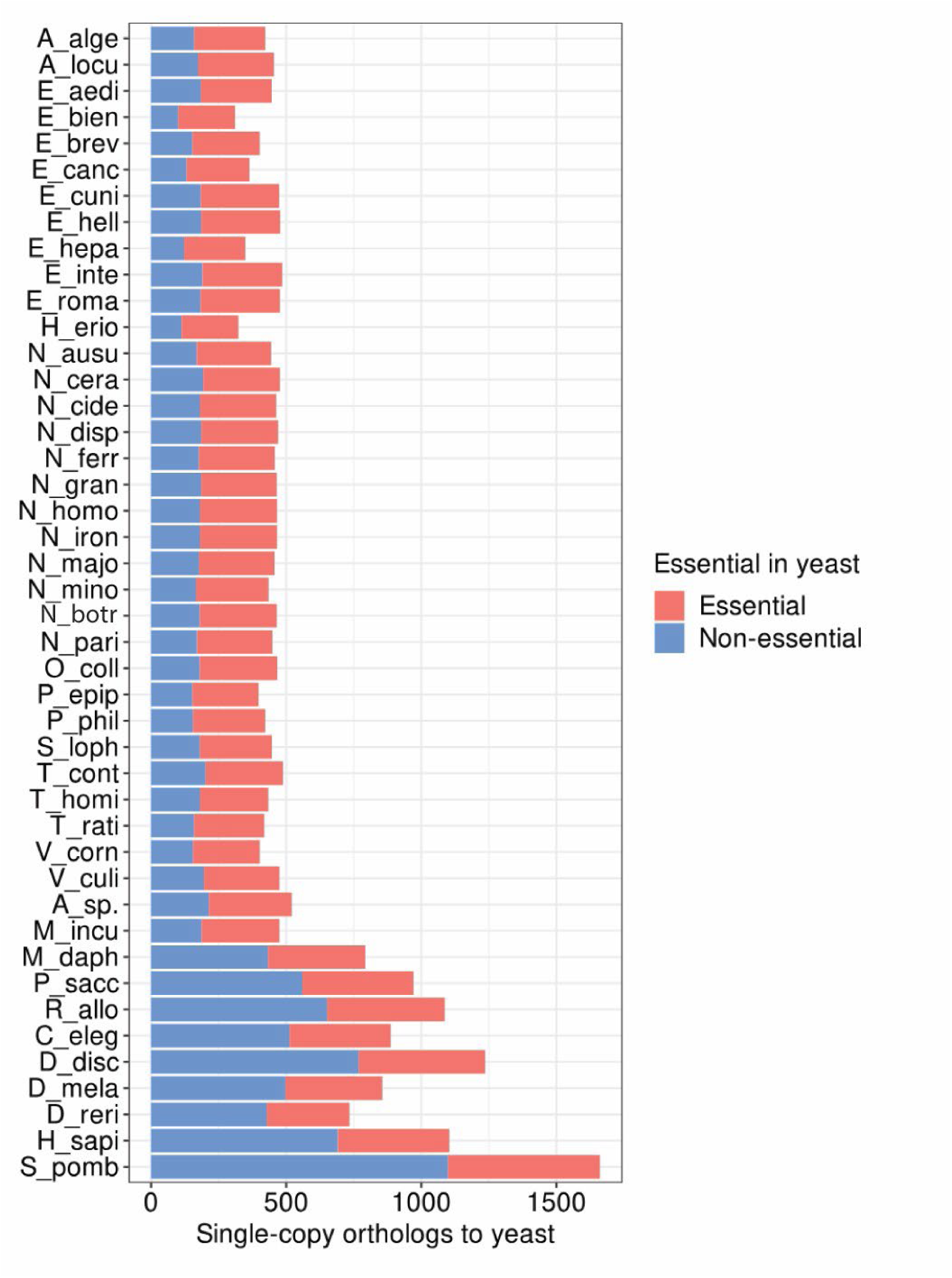
Number of single-copy orthologs from each species that are essential or non-essential in yeast. The number of single-copy orthologs that each species has compared to yeast is shown, with orthologs of essential and non-essential genes in yeast shown according to the legend at the right.

**Supplementary fig. S2.**
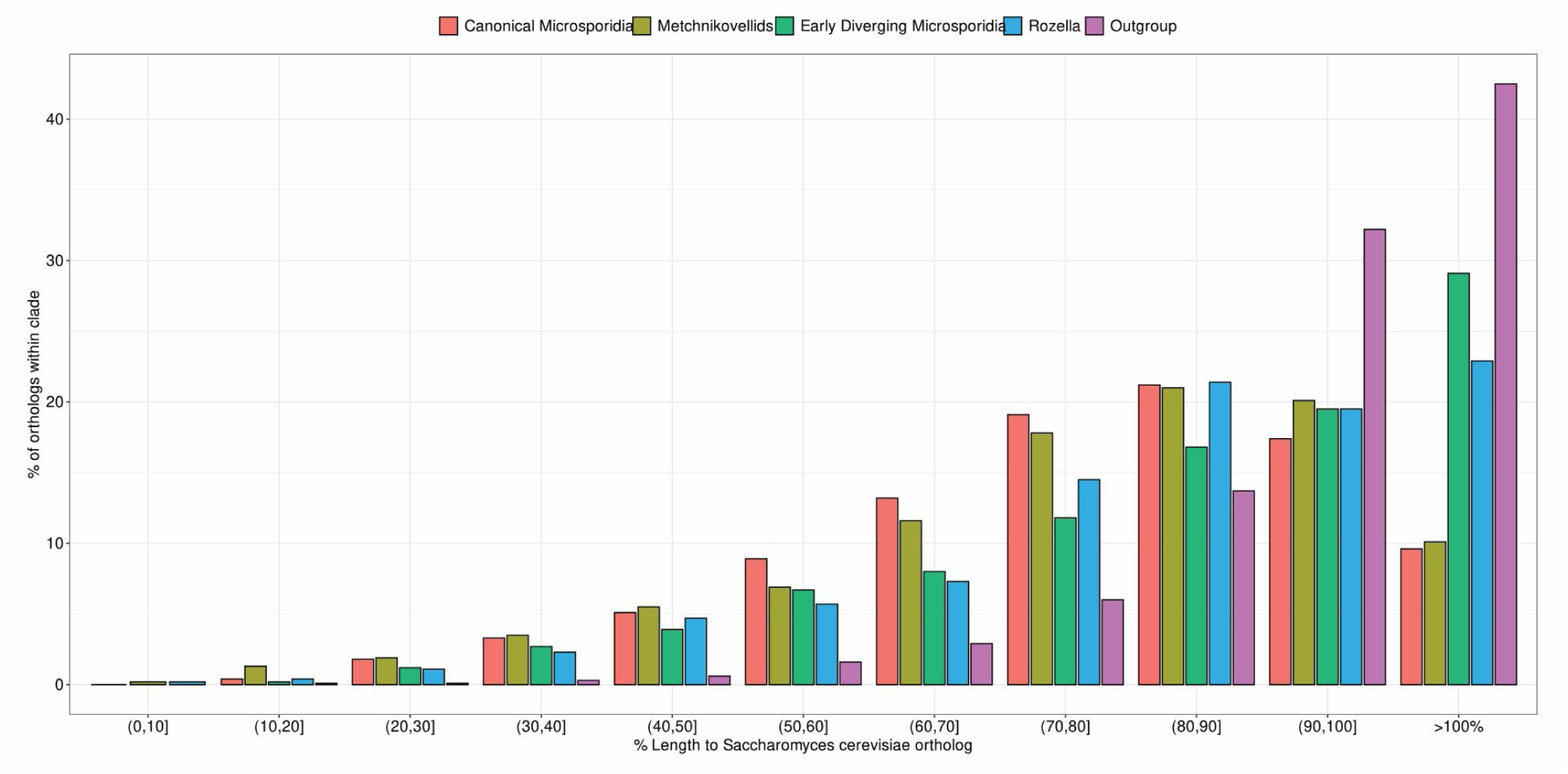
Length distribution of orthologs compared to yeast. Distribution of the relative ortholog lengths relative to yeast for each species group.

**Supplementary fig. S3.**
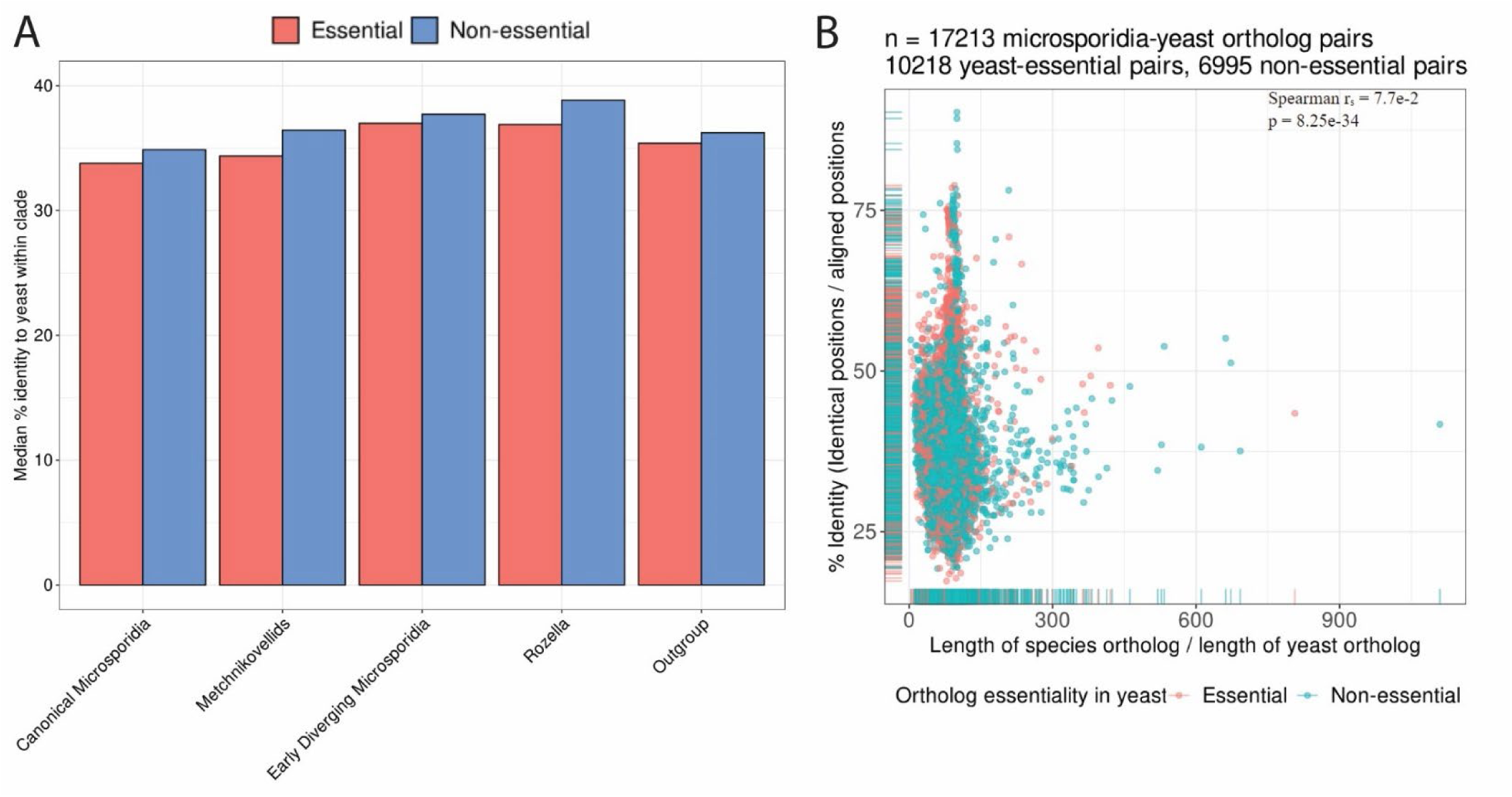
Sequence identity of orthologs. (A) Distribution of sequence identity relative to yeast ortholog of the indicated groups of species stratified by yeast-essential and non-essential orthologs. (B) Relative lengths of microsporidia orthologs to yeast compared to their percent identities.

**Supplementary fig. S4.**
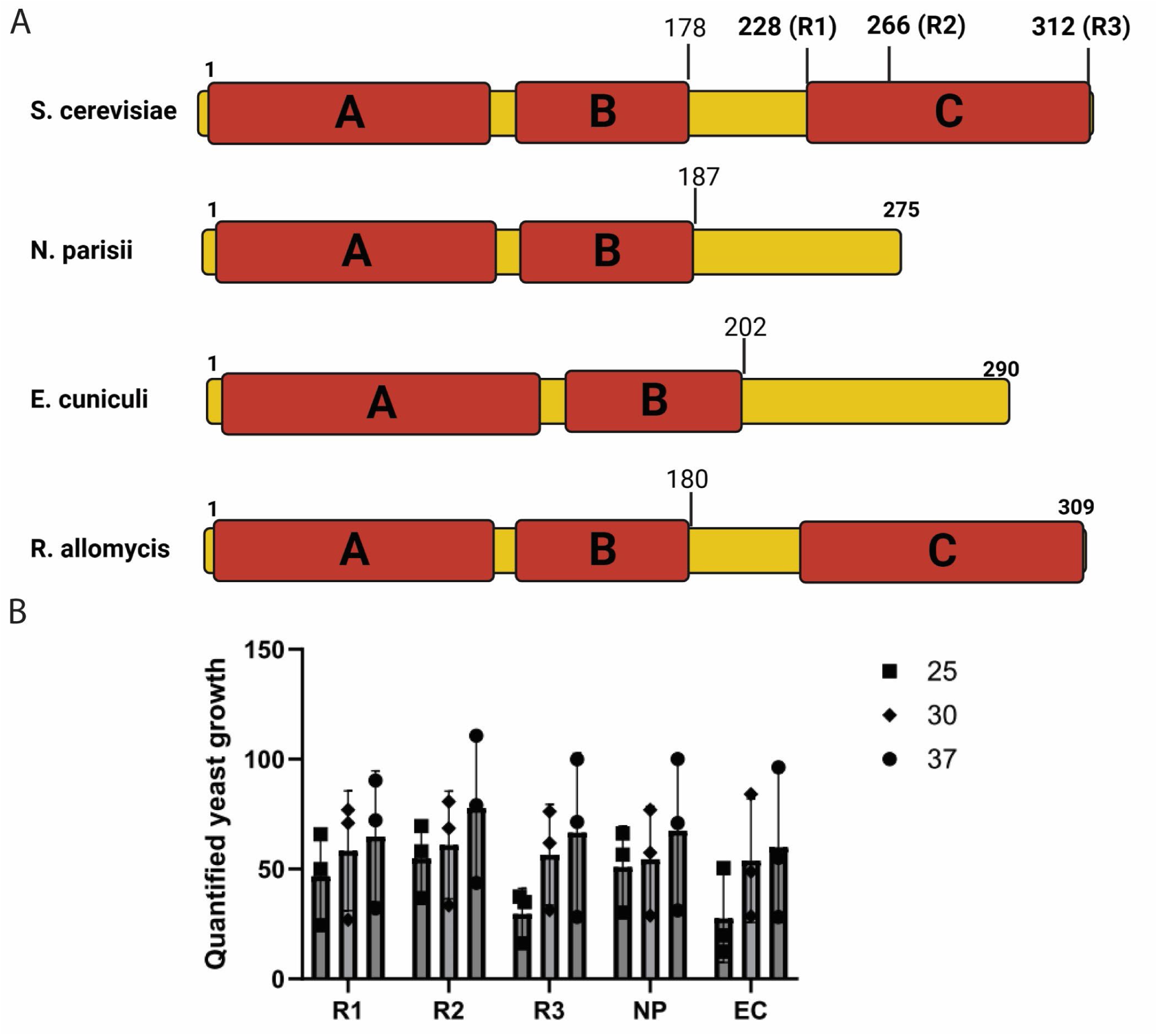
*S. cerevisiae* RPPO truncations rescue to similar extent as microsporidia orthologs. (A) Domain diagram of *S. cerevisiae* and microsporidia RPP0 proteins. Domain identities as given: A (Large ribosomal subunit protein uL10), B (Large ribosomal subunit protein uL10-like, insertion domain), C (60s Acidic ribosomal protein). (B) Various RPP0 containing-plasmids were transformed into a hetKO deletion of RPPO. These strains were dilution plated at the indicated temperature and colony size was quantified. All statistical comparisons between strains at each temperature have p-values > 0.5 determined using Two-way ANOVA.

**Supplementary fig. S5.**
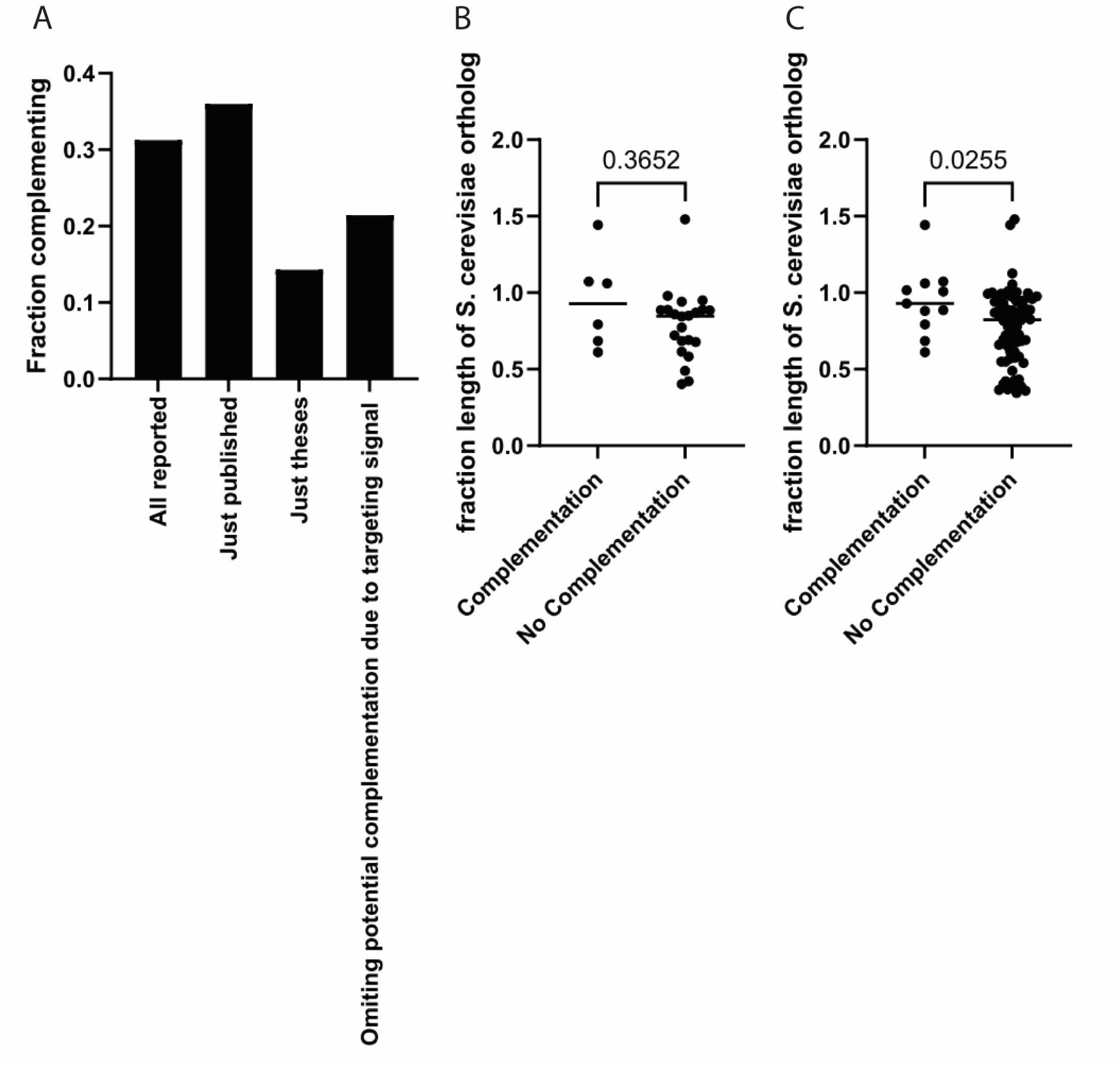
Previously reported complementation by microsporidia orthologs. (A) Fraction of microsporidia proteins that complement deletion strains of their yeast orthologs. (B-C) Ratio of microsporidia/ yeast length for each protein that could complement or not complement. (B) Literature reported complementation, omitting potential complementation due to targeting signals. (C) Combined literature complementation (fig. S5B) and complementation from this study (fig. 6C). The p-values were determined by Mann-Whitney test.

